# Vertical Ground Reaction Force Morphology Is Determined by Step-to-Step Transition Mechanical Energy Imbalance During Human Walking

**DOI:** 10.64898/2026.03.09.710627

**Authors:** Seyed-Saleh Hosseini-Yazdi, John EA Bertram

**Author notes:** **Correspondence Address:** John EA Bertram, Biomedical Engineering Department, Schulich School of Engineering, University of Calgary, 2500 University Dr NW, Calgary AB T2N1N4.

## Abstract

Vertical ground reaction force (vGRF) profiles during walking typically exhibit a double-peaked structure with a mid-stance trough, yet the mechanical conditions governing this morphology remain incompletely defined. In this study, we examined how the balance between push-off and collision impulses during the step-to-step transition influences the temporal and structural characteristics of the vGRF trajectory. Empirical relationships describing push-off and collision work were used to compute transition impulses across walking speeds ranging from 0.8 to 1.4 m·s□^1^. A normalized Impulse Balance Index (IBI) was defined to quantify the relative dominance of push-off and collision impulses. The temporal position of the mid-stance trough was quantified using a Trough Deficit Index (TDI) derived from quadratic fits of the vGRF trajectory. Across walking speeds, push-off and collision variations produced step-to-step active work performance imbalance. Push-off and collision became approximately balanced near 1.2 m·s□^1^, corresponding to the mechanically preferred walking speed. Deviations from this balanced condition were associated with systematic shifts in trough timing: the trough occurred 1.83% and 1.56% earlier in stance at 0.8 and 1.0 m·s□^1^, respectively, and 1.31% later at 1.4 m·s□^1^ relative to the reference speed. TDI exhibited a strong inverse relationship with impulse balance (IBI), indicating that vGRF morphology is tightly coupled to the mechanical balance of the step transition. A simplified pendular model further demonstrated that active torque, representing work, during single support shifts the quadratic vertex of the force trajectory by approximately 48.6–51.1% of stance, consistent with the observed trough timing variations. These results show that vertical GRF morphology reflects the imbalance between push-off and collision provides a simple signal of step-to-step transition mechanics, that may be used for rehabilitation and exoskeleton modulation.

## Introduction

Human walking is achieved by applying forces against the ground to accelerate and redirect the body’s center of mass (COM) [1]. These forces, measured as ground reaction forces (GRFs), provide direct insight into the mechanical work performed by the lower limbs during locomotion [2]. Among the three components of GRF, the vertical component is the largest because it must support body weight and regulate vertical motion of the COM throughout stance [3]. Consequently, vertical ground reaction force (vGRF) profiles have been widely used to characterize human gait mechanics and evaluate locomotor function [4]. Because GRFs directly reflect the mechanical interaction between the body and the ground, they also provide a potentially useful signal for wearable and soft robotic devices to detect propulsion deficits and adaptively assist gait during rehabilitation.

During normal walking, the vGRF trajectory typically exhibits a characteristic double-peaked pattern within the stance phase [5]. The first peak occurs shortly after heel strike and is associated with weight acceptance [6] as the leading limb decelerates the downward and forward motion of the COM. The second peak occurs near toe-off and reflects propulsion generated primarily by the ankle plantarflexors to accelerate the body forward and upward into the next step [6–8]. Between these peaks, a trough occurs during mid-stance when the body vaults over the stance limb and the COM reaches its highest position [6]. This double-hump pattern has been consistently observed across a wide range of walking speeds and populations and is considered a fundamental feature of human gait dynamics.

The mechanical interpretation of these force features has been closely linked to the energetics of the step-to-step transition [2]. During walking, the COM velocity vector must be redirected from one stance limb to the next [9]. This transition involves a coordinated sequence of mechanical work performed by the trailing and leading limbs [10]. Preemptive push-off generated by the trailing limb adds positive mechanical work that helps redirect the COM velocity before heel strike. Immediately after, the leading limb performs negative work through collision as the new stance limb accepts the body weight. This push-off–collision sequence reduces energy losses and is a major determinant of the mechanical and metabolic cost of walking [9].

Simplified models of walking have proven useful in explaining these energetic interactions [11– 13]. The “simplest walking model,” which represents the body as a point mass supported by rigid legs, predicts that optimal walking occurs when push-off work approximately compensates for collision losses during the step-to-step transition [9]. Under this condition, the mechanical energy of the COM remains nearly constant and the subsequent single-support phase can proceed passively as an inverted pendulum [9,14,15]. Consistent with this concept, classical inverted pendulum models describe the stance phase as a passive vaulting motion in which the COM rises and falls while the body rotates over the stance limb [16]. These models predict a symmetric force trajectory during single support, with the minimum vertical force occurring near mid-stance when the COM passes over the stance foot [9].

Despite these theoretical insights, vGRF profiles observed experimentally are not always symmetric [17,18]. Changes in walking speed, terrain conditions, neuromotor control, or pathological gait can alter the relative magnitudes of the two vertical force peaks. For example, increases in walking speed are associated with systematic changes in peak forces and impulses, indicating that the mechanical balance between push-off and collision varies with locomotor conditions [19]. Similarly, alterations in musculoskeletal function or assistive devices can modify the distribution of vertical forces during weight acceptance and late stance [6]. These observations suggest that deviations from symmetric vGRF trajectories may reflect changes in how mechanical work is distributed across the stance phase.

However, the mechanical conditions that determine whether the vGRF profile becomes symmetric or asymmetric remain insufficiently defined. In particular, it is not fully understood how the relative magnitudes of collision and push-off impulses influence the timing of the mid-stance trough and the resulting skewness of the vertical force trajectory. Clarifying this relationship would provide a simple means of interpreting vGRF morphology in terms of underlying energetic balance during walking.

In this study, we examine how the balance between collision and push-off work influences the vertical GRF trajectory during walking. First, we use a simplified step-to-step transition model to establish the relationship between transition impulses and the amplitudes of the two vertical force peaks. Second, we analyze single-support dynamics using a pendular model to determine how active torque alters the symmetry of the vertical force trajectory. Finally, we compare these model predictions with empirical observations of walking across different speeds. Based on this framework, we introduce a simple metric—the Vertical GRF Trough Timing Index (vGRF-TTI)—to quantify the temporal position of the mid-stance trough and interpret it as an indicator of the balance between push-off and collision work during human walking. Such a mechanically interpretable signal may also provide a practical control variable for wearable robotic systems to detect propulsion deficits and modulate assistive actuation during gait rehabilitation.

## Materials and Methods

Human walking requires periodic redirection of the center-of-mass (COM) velocity during the transition from one stance limb to the next. This redirection occurs through a coordinated sequence of mechanical work performed by the trailing and leading limbs. During late stance, the trailing limb generates push-off work, which accelerates the COM upward and forward. Immediately following heel strike, the leading limb performs collision work, which decelerates the COM as the body is redirected onto the new stance limb.

The energetic balance between push-off and collision determines the mechanical efficiency of the step-to-step transition. When push-off work approximately compensates for collision losses, the mechanical energy of the COM remains nearly constant across steps, allowing the subsequent single-support phase to proceed with minimal active regulation. Deviations from this balance indicate either insufficient propulsion or excess energy input during the transition.

To examine how this balance varies with walking speed, empirical relationships describing push-off and collision work as functions of speed are used (0.8 m.s^-1^ to 1.4 m.s^-1^). These relationships were derived from previously reported measurements of COM work trajectories during human walking [20]. Push-off work *w*_*po*_and collision work *w*_*co*_are expressed as polynomial functions of walking speed. From these relationships, the corresponding transition impulses are estimated as [21]:

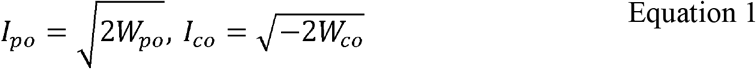

which provide approximations of the vertical impulses associated with push-off and collision during the double-support phase (Figure 1A).

**Figure 1.**
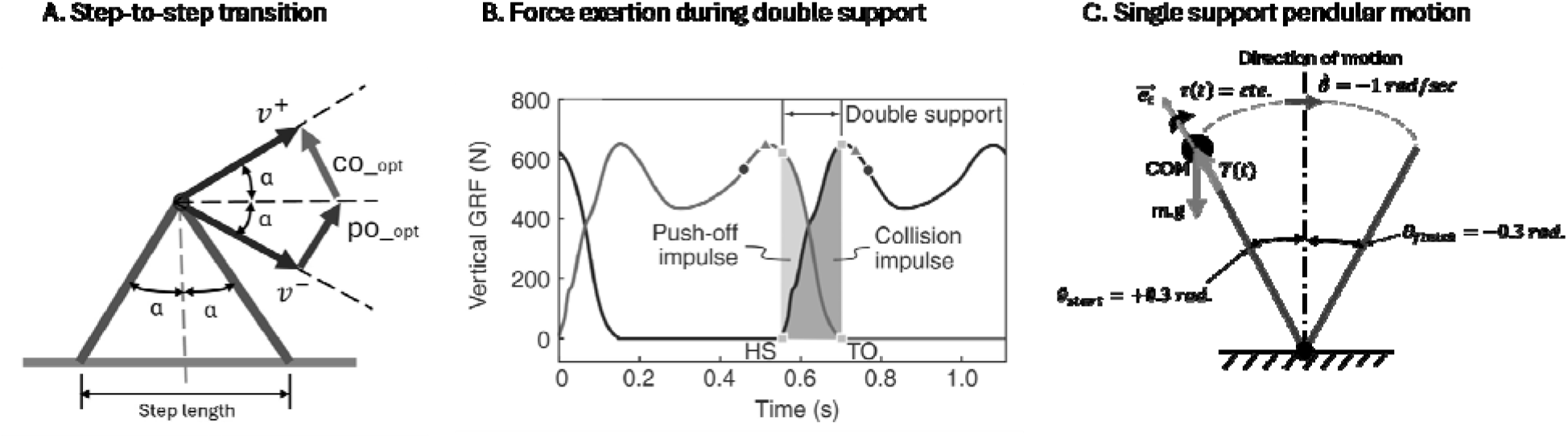
(A) the simple model of step-to-step transition in which transition is performed by push-off followed by a collision impulse, (B) human walking push-off and collision performance based on GRF (adopted from *[1]*), and (C) the pendular motion of the single support to complete the stance.

To quantify the balance between push-off and collision impulses, an **Impulse Balance Index (IBI)** is defined as:

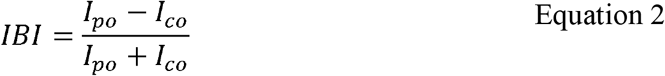

where *I*_*po*_and *I*_*co*_denote the push-off and collision impulses, respectively. This normalized index ranges between −1 and 1 and provides a dimensionless measure of transition balance.

An index value near zero indicates approximately balanced impulses, whereas positive or negative values indicate dominance of push-off or collision impulses, respectively. Because human walking typically exhibits mechanical equilibrium near the preferred walking speed[13], the index is adjusted relative to the reference speed of 1.2 m.s^-1^, such that *IBI*^∗^ = *IBI − IBI*_*ref*_, where *IBI*_*ref*_ is the value computed at the preferred walking speed. This adjustment allows deviations from balanced transition mechanics to be evaluated across different walking speeds.

The collision peak and push-off peak of the vGRF trajectory are detected using a peak-finding algorithm based on prominence and minimum separation criteria. Collision impulse is estimated as the area under the force curve from heel-strike to the first peak, while push-off impulse is estimated as the area from the second peak to toe-off [1] (Figure 1B).

The dynamics of the center of mass during single support are represented using a simplified inverted pendulum model. In this model, the body mass is concentrated at the pelvis and supported by a rigid stance limb of length *l*. The motion of the COM follows pendular dynamics governed by gravity and centripetal acceleration. The axial force along the stance limb is expressed as:

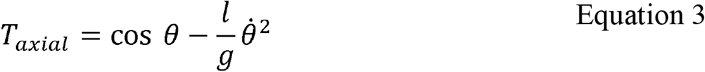

where *θ* is the stance-leg angle and 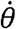 is the angular velocity. The vertical component of the ground reaction force is obtained by projecting this axial force onto the vertical direction: *F*_*z*_ =*T*_*axial*_ cos *θ*. To examine the effect of active mechanical energy regulation during single support, constant hip torques are introduced into the pendular dynamics:

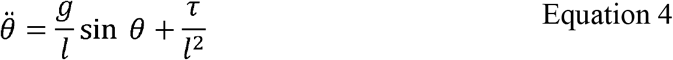

where *τ* represents the applied hip torque. Simulations are initiated at a stance-leg angle of 0.3 rad with initial angular velocity −1.1 rad.s^-1^ and terminated when the stance leg reached the symmetric terminal angle of −0.3 rad (Figure 1C).

To quantify deviations of empirical force trajectories from ideal pendular mechanics, a **Pendular Deviation Index (PDI)** is defined. For each stance phase, the empirical vGRF trajectory *F*_*z*_ (*t*) is compared with its quadratic approximation *F*_*z,quad*_ (*t*). The deviation index is computed as:

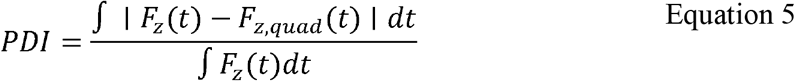

This normalized measure represents the relative magnitude of deviations from quadratic pendular behavior. To evaluate changes in pendular deviation across walking speeds, the index is referenced to the mechanically preferred walking speed[13] (1.2 m.s^-1^): *PDI*^∗^ = *PDI* − *PDI*_*ref*_,where *PDI*_*ref*_ is the value obtained at the preferred speed. Positive values therefore indicate greater deviation from passive pendular mechanics relative to the reference condition.

To quantify deviations in the temporal structure of the vertical ground reaction force trajectory, a **Trough Deficit Index (TDI)** is defined based on the timing of the mid-stance minimum of the vertical ground reaction force (vGRF). During walking, the vGRF typically exhibits a double-peaked pattern consisting of an early collision peak and a late push-off peak, separated by a trough occurring during mid-stance when the center of mass vaults over the stance limb. The vGRF trajectory during stance is approximated using a quadratic function: . where represents normalized stance time (% stance). The timing of the trough is determined analytically from the vertex of the quadratic function: –.

where denotes the temporal location of the minimum vGRF during stance. Because balanced push-off and collision impulses occur near the preferred walking speed, the trough timing at 1.2 m·s□^1^ was used as a reference condition. The **Trough Deficit Index** is therefore defined as:

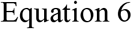

where is the trough timing at the preferred walking speed and is the trough timing observed at each walking condition. Under this definition, corresponds to the mechanically balanced walking condition, whereas positive values indicate an earlier occurrence of the mid-stance trough relative to the reference condition. The TDI therefore provides a simple morphology-based indicator of deviations from balanced step-to-step transition mechanics.

## Results

Empirical analysis of step-to-step transition mechanics revealed systematic changes in push-off and collision work across walking speeds. As walking speed increased from 0.8 to 1.4 m·s□^1^, push-off work increased from approximately 0.16 J·kg□^1^ to 0.38 J·kg□^1^, while the magnitude of collision dissipation varied from 0.06 J·kg□^1^ to 0.44 J·kg□^1^. The corresponding push-off and collision impulses ranged from 0.41 to 0.61 m·s□^1^ and 0.24 to 0.67 m·s□^1^, respectively (Figure 2).

**Figure 2.**
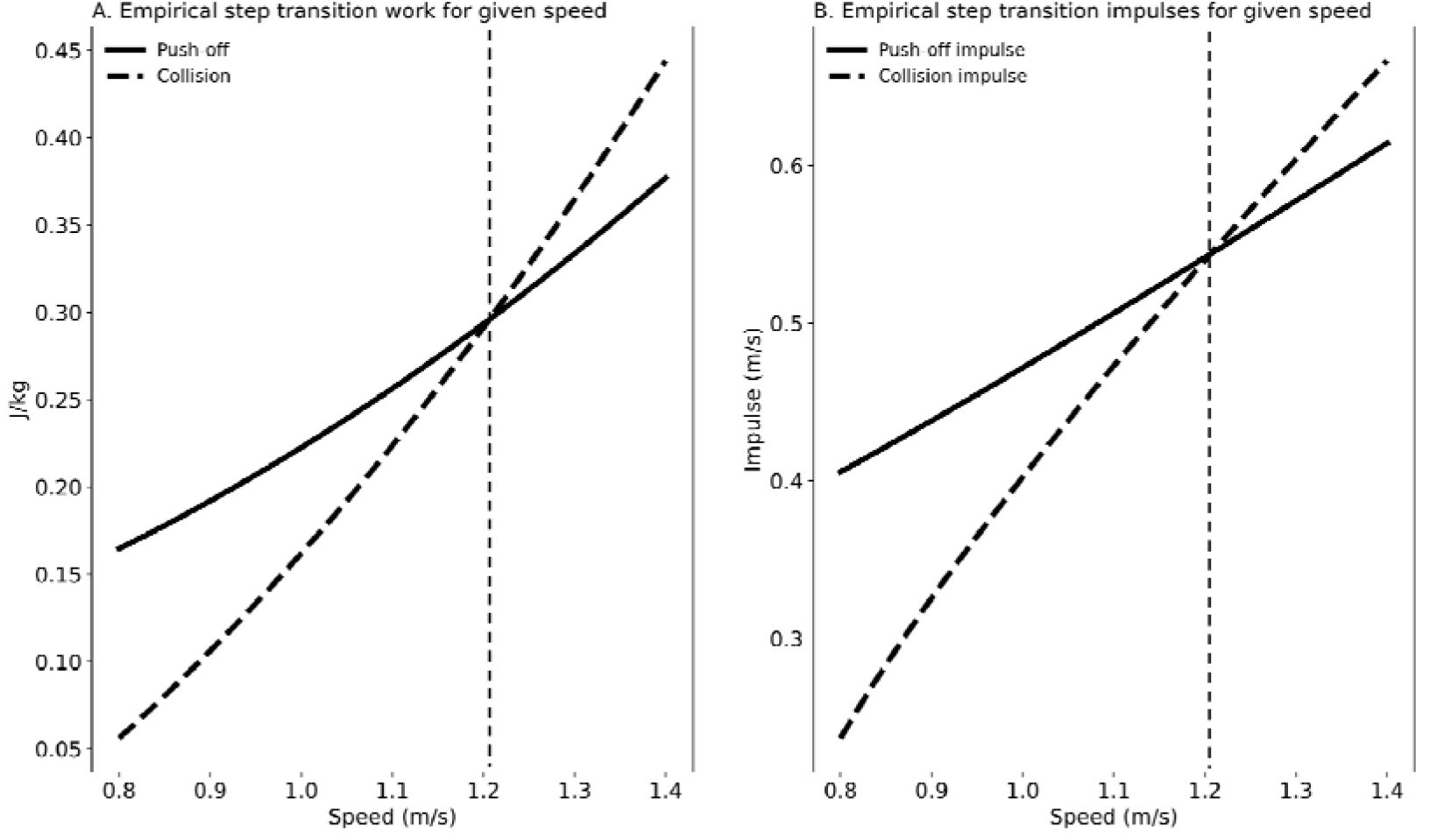
(A) Empirical push-off and collision work trajectories as functions of walking speed. (B) Corresponding push-off and collision impulses derived from the work trajectories. There is only one speed for which the push-off and collision are balanced.

At lower walking speeds, push-off impulses exceeded those associated with heel-strike collisions, whereas at higher speeds the opposite trend was observed, with collision impulses becoming larger than push-off impulses. The vertical GRF peaks were therefore generally asymmetric across walking speeds. At approximately 1.21 m·s□^1^, push-off and collision impulses were nearly equal, indicating a balanced step-to-step transition.

Changes in transition balance were accompanied by systematic shifts in the timing of the mid-stance trough of the vertical GRF trajectory. The Impulse Balance Index (IBI), evaluated relative to the reference condition at 1.2 m·s□^1^, ranged from 0.0617 at 0.8 m·s□^1^ to −0.0114 at 1.4 m·s□^1^, with the reference condition yielding IBI = 0. The IBI decreased linearly with walking speed (slope = −0.12,) significantly (, Figure 3).

**Figure 3.**
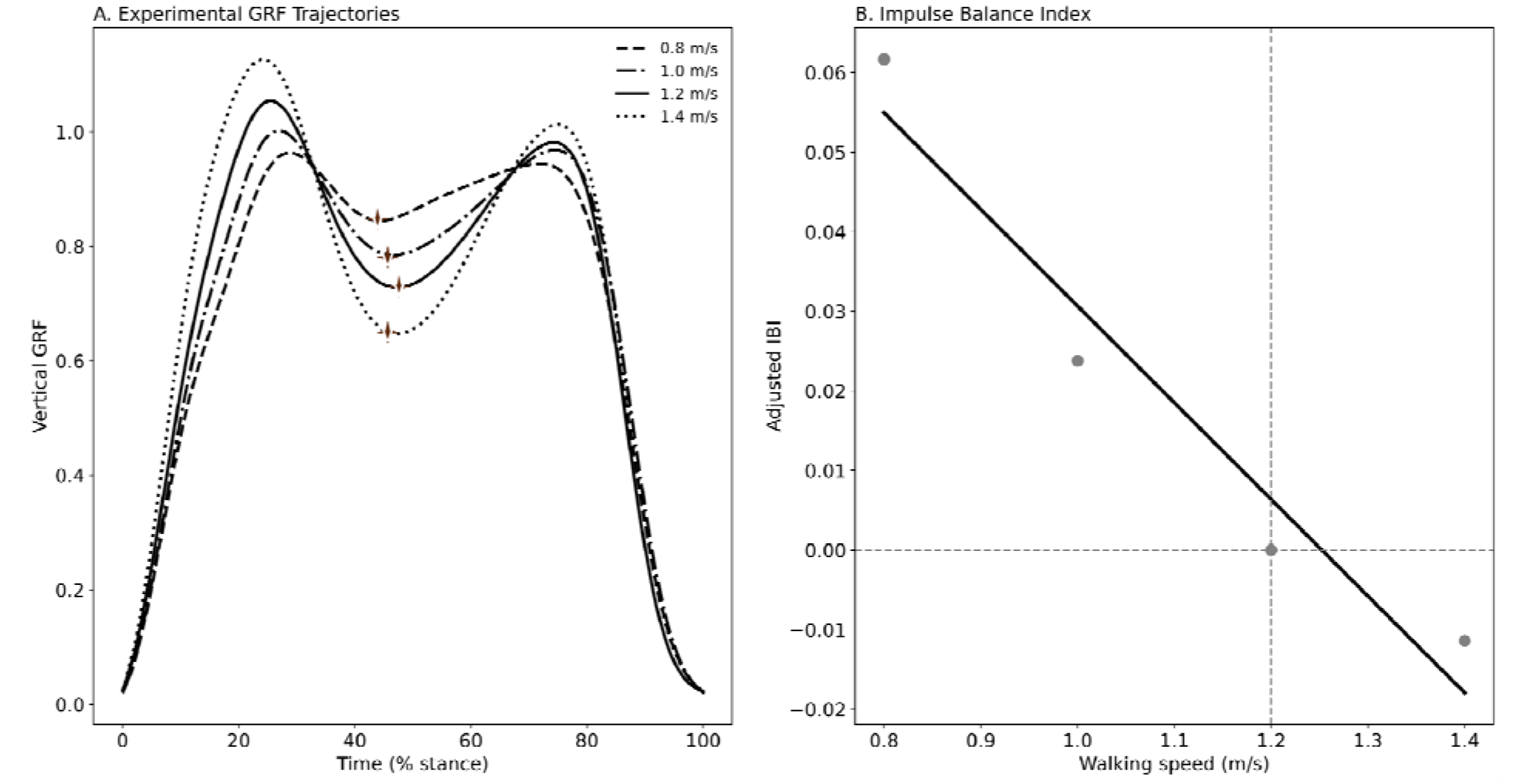
(A) Vertical ground reaction force profiles during stance for different walking speeds (adopted from *[20]*). The midstance troughs are indicated by starts. (B) Adjusted Impulse Balance Index (IBI) as a function of walking speed that depicts an inverse relationship.

The Trough Deficit Index (TDI) quantified the shift in trough timing relative to this reference condition. At 0.8 and 1.0 m·s□^1^, the trough occurred earlier in stance with values of −1.83% and −1.56% of stance, respectively, whereas at 1.4 m·s□^1^ the trough occurred later by 1.31% of stance. The Trough Deficit Index (TDI) increased significantly with walking speed (slope = 5.49,,). Across walking speeds, trough timing was strongly associated with impulse balance. Linear regression between TDI and IBI yielded a correlation coefficient of with a slope of −40.4 (Figure 4).

**Figure 4.**
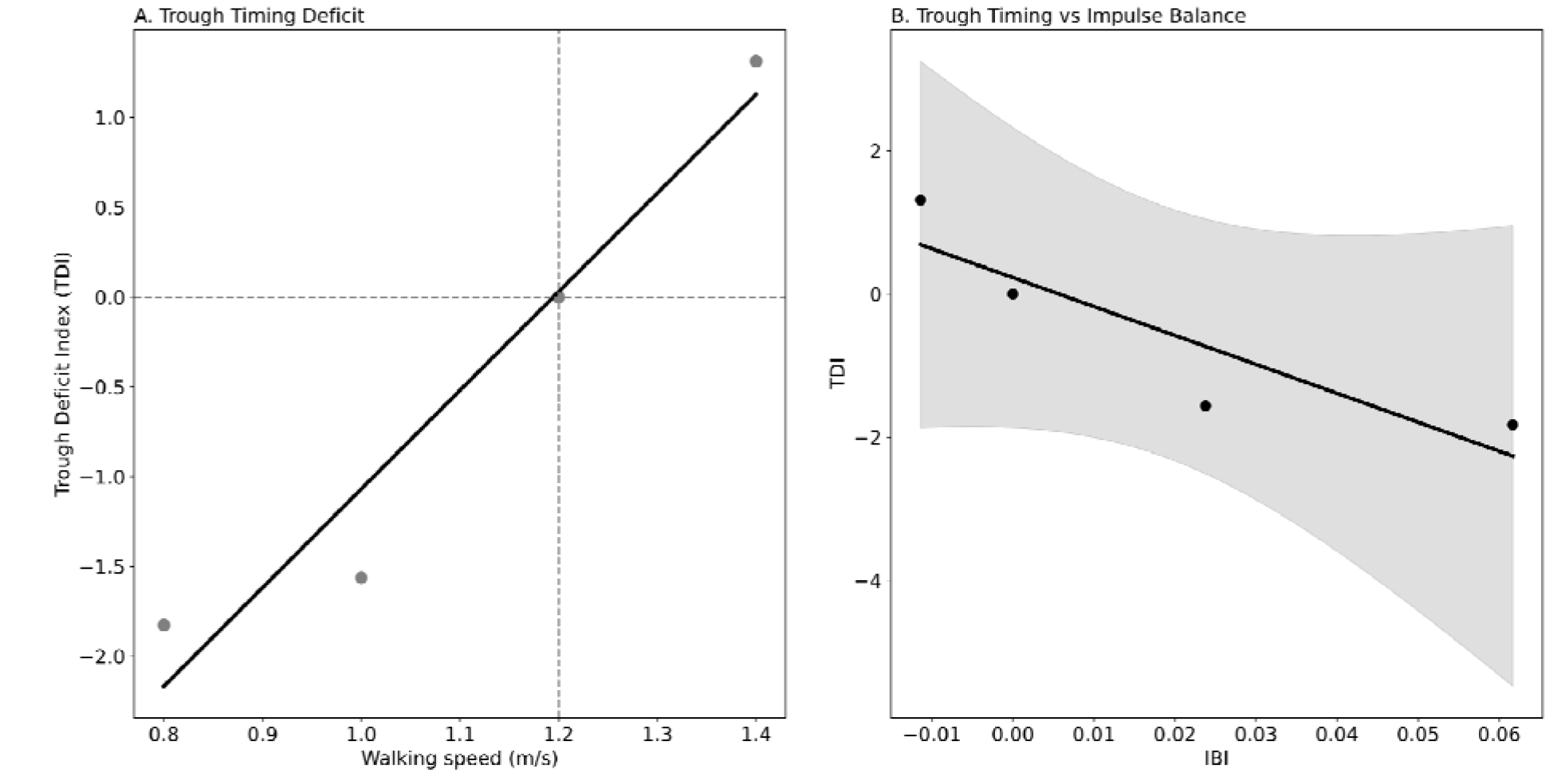
(A) Trough Deficit Index (TDI), representing the shift in mid-stance force trough timing relative to the reference speed (1.2 m·s□^1^), increases with walking speed. (B) TDI shows an inverse linear relationship with the Impulse Balance Index (IBI), indicating that changes in collision–push-off impulse balance are associated with systematic shifts in GRF trough timing.

Simulations using the quadratic pendular model were used to examine how applied hip torque influenced the symmetry of the vertical GRF trajectory during single support. The predicted timing of the quadratic vertex ranged between 48.6% and 51.1% of stance depending on the applied torque magnitude (Figure 5A).

**Figure 5.**
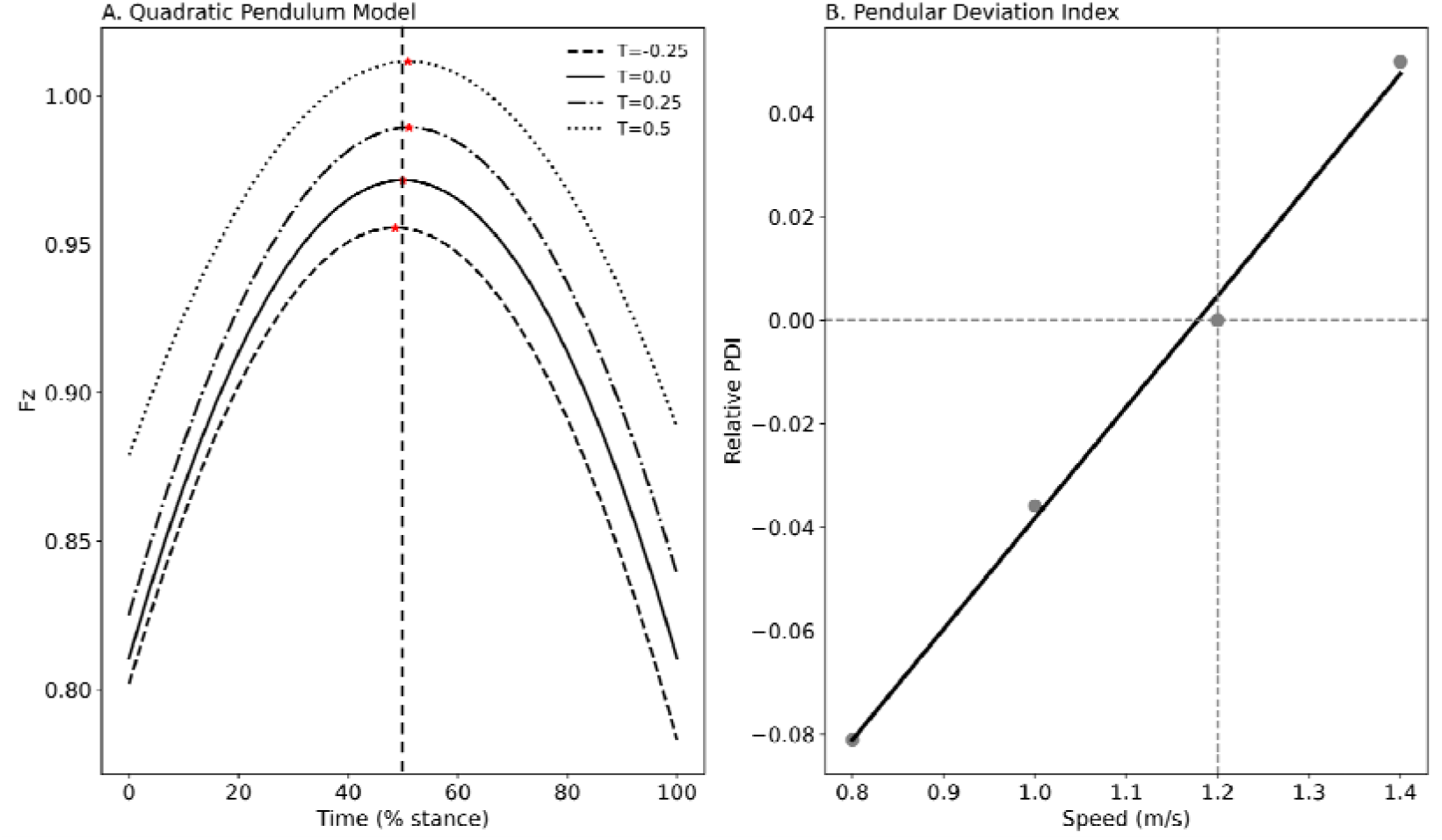
(A) Quadratic inverted pendulum model predictions of vertical ground reaction force for different constant torques ‘’ representing addition or dissipation of the COM mechanical energy. The stars indicate the mid-stance force trough. (B) Variation of Relative Pendular Deviation Index (PDI) derived from experimental GRF across walking speeds indicates a direction linear relationship.

The Pendular Deviation Index (PDI) was used to quantify deviations of empirical vertical GRF trajectories from quadratic pendular behavior. Relative to the reference walking speed of 1.2 m·s□^1^, PDI values were −0.081 and −0.036 at 0.8 and 1.0 m·s□^1^, respectively, and 0.050 at 1.4 m·s□^1^. Relative PDI increased linearly with walking speed (slope = 0.214,) significantly (, Figure 5B).

## Discussion

The present study examines how the balance between push-off and collision impulses influences the morphology of the vertical ground reaction force (vGRF) trajectory during walking. The results demonstrate that changes in step-to-step transition mechanics are reflected not only in the magnitudes of vertical impulses but also in the temporal structure of the vGRF trajectory. In particular, deviations from balanced push-off and collision impulses are accompanied by systematic shifts in the timing of the mid-stance trough. These findings indicate that the temporal position of the trough provides a simple observable indicator of the mechanical balance governing the step-to-step transition.

The energetic basis of this relationship arises from the mechanics of redirecting the center-of-mass (COM) velocity during walking [1]. During each step transition, the trailing limb performs positive work through push-off while the leading limb performs negative work during collision at heel strike [2]. When push-off work approximately compensates for collision losses, the mechanical energy of the COM remains nearly constant across steps, allowing the subsequent single-support phase to proceed with minimal active intervention [9,10]. This principle is a central prediction of simple walking models and has been widely used to explain the energetic cost of walking [2,16]. The empirical results obtained here are consistent with this framework, showing that push-off and collision impulses become nearly equal near the preferred walking speed of approximately 1.2 m·s□^1^ [13].

These findings also align with classical descriptions of walking mechanics in which the center of mass vaults over the stance limb in an inverted-pendulum-like motion [14,15,22,23]. In such models, the vertical force reaches a minimum when the COM passes directly above the stance foot [24]. However, deviations from symmetric force trajectories arise when the balance between push-off and collision work is altered. The present results suggest that these deviations manifest as measurable shifts in trough timing within the vGRF trajectory.

Beyond confirming the energetic balance predicted by transition models, the present study demonstrates that trough timing is strongly associated with impulse balance. The Trough Deficit Index (TDI) varied systematically across walking speeds and showed a strong inverse correlation with the Impulse Balance Index (IBI). This relationship indicates that the temporal structure of the vGRF trajectory reflects the underlying balance between push-off and collision impulses.

The trough of the vertical GRF occurs when the COM reaches its highest position during stance. Because this event marks the transition between early-stance loading and late-stance propulsion, its timing is inherently influenced by how mechanical work is distributed between the two limbs. Previous studies have shown that variations in walking speed and propulsion alter the distribution of mechanical work during stance [2,16,25]. The present results extend these findings by demonstrating that such variations are encoded directly in the morphology of the vertical GRF trajectory.

An important implication of this result is that trough timing can serve as a practical indicator of transition mechanics without requiring explicit calculation of work or impulse integrals. Because the trough can be detected directly from the force trajectory, this metric provides a simple signal that can be extracted from vertical force measurements alone. Such a feature may be particularly useful in wearable sensing systems or instrumented insoles where direct measurement of full-body energetics is not feasible.

The Pendular Deviation Index (PDI) is introduced to quantify deviations of empirical vGRF trajectories from quadratic pendular behavior. The results indicate that walking at lower speeds more closely resembled passive pendular dynamics, whereas deviations from pendular behavior increased at higher speeds. These findings are consistent with previous observations that the inverted pendulum model becomes less accurate as walking speed increases and active muscle work becomes more prominent [2,16,25,26].

We suggest that the deviations from pendular behavior observed in the vertical ground reaction force trajectories are closely associated with the relative imbalance between push-off and collision impulses. As walking speed deviated from the condition where push-off and collision are balanced, the distribution of vertical forces during stance became increasingly asymmetric. This redistribution of forces altered the morphology of the GRF trajectory, shifting the relative magnitudes of the two peaks and modifying the timing of the mid-stance trough, necessitating an active positive work performance during the single support [3,17,27]. The increase in the Pendular Deviation Index (PDI) therefore reflects not simply an increase in active work with speed, but a change in how mechanical work is partitioned between the trailing and leading limbs during the step-to-step transition and subsequent ingle support.

Simulations using the quadratic pendular model provided a mechanistic framework for interpreting these empirical observations [14,15]. The model demonstrated that applying hip torques during single support, representing single support active work performance, systematically shifts the timing of the quadratic vertex of the vertical force trajectory. Although the simplified pendular model does not capture the full complexity of human locomotion, it provides a useful conceptual framework for understanding how active mechanical work influences force trajectories. The model results show that relatively small changes in applied torque (single support active work) can produce measurable shifts in trough timing, supporting the interpretation that variations in GRF morphology reflect changes in mechanical regulation during stance.

The identification of simple GRF-based indices that reflect transition mechanics has potential implications for gait monitoring and assistive device control. Vertical ground reaction forces can be measured using instrumented treadmills [28], force plates [22], wearable insoles [6], or soft robotic sensors. Indices such as **IBI, TDI**, and **PDI** could therefore serve as practical signals for detecting changes in locomotor mechanics using force measurements alone. In rehabilitation contexts, propulsion deficits [29] are a common feature of impaired gait [8], particularly in populations such as individuals recovering from stroke [30]. Reduced push-off from the paretic limb alters the balance of transition impulses and can increase the mechanical cost of walking [17]. Metrics derived from vertical GRF morphology may therefore provide a useful means of identifying propulsion deficits and monitoring rehabilitation progress. In addition, assistive robotic devices could potentially use such simple signals to detect transition imbalance and adaptively modulate assistance during walking.

Several limitations should be considered when interpreting the present results. First, the empirical relationships describing push-off and collision work are derived from previously reported data. Although these relationships capture well-established trends in locomotion energetics, individual variability in gait mechanics may influence the precise balance of impulses. Second, the quadratic approximation used to characterize the vGRF trajectory provides a simplified representation of the force profile. While this approach allows analytical estimation of trough timing and facilitates comparison with pendular models, the actual force trajectory during walking is influenced by complex musculoskeletal dynamics that are not fully captured by a quadratic representation [31]. Finally, the simplified pendular model used in this study represents the body as a point mass supported by a rigid leg and therefore does not include multi-joint muscle coordination [31], compliant tissues [32], or neuromotor control mechanisms [33]. As a result, the model should be interpreted primarily as a conceptual tool for understanding how active mechanical work may influence force symmetry rather than as a detailed representation of human gait dynamics.

The present study demonstrates that the morphology of the vertical ground reaction force trajectory reflects the mechanical balance governing the step-to-step transition during walking. Empirical analysis showed that push-off and collision impulses become balanced near the preferred walking speed, and that deviations from this balance are accompanied by systematic shifts in the timing of the mid-stance trough. The strong relationship between trough timing and impulse balance indicates that the temporal structure of the vertical GRF trajectory provides a simple observable signal of transition mechanics. Together with the pendular model results, these findings suggest that vertical GRF morphology encodes information about both the energetic balance of the step transition and the active regulation of single-support dynamics.

## References

[1] P.G. Adamczyk, A.D. Kuo, Redirection of center-of-mass velocity during the step-to-step transition of human walking, J Exp Biol 212 (2009) 2668–2678. 10.1242/jeb.027581.

[2] J.M. Donelan, R. Kram, A.D. Kuo, Mechanical work for step-to-step transitions is a major determinant of the metabolic cost of human walking, J Exp Biol 205 (2002) 3717–3727. 10.1242/jeb.205.23.3717.

[3] S. Apte, M. Plooij, H. Vallery, Simulation of human gait with body weight support: benchmarking models and unloading strategies, J NeuroEngineering Rehabil 17 (2020) 81. 10.1186/s12984-020-00697-z.

[4] E. Chung, S.-H. Lee, H.-J. Lee, Y.-H. Kim, Comparative study of young-old and old-old people using functional evaluation, gait characteristics, and cardiopulmonary metabolic energy consumption, BMC Geriatr 23 (2023) 400. 10.1186/s12877-023-04088-6.

[5] A.M.F. Barela, P.B. de Freitas, M.L. Celestino, M.R. Camargo, J.A. Barela, Ground reaction forces during level ground walking with body weight unloading, Braz J Phys Ther 18 (2014) 572–579. 10.1590/bjpt-rbf.2014.0058.

[6] T.L. Libera, J. Streamer, R.M. Queen, Peak Weight Acceptance, Mid Stance Trough, and Peak Push-Off Force Symmetry Are Decreased in Older Adults Compared With Young Adults, Journal of Applied Biomechanics 41 (2025) 117–123. 10.1123/jab.2024-0027.

[7] M.G. Browne, J.R. Franz, More push from your push-off: Joint-level modifications to modulate propulsive forces in old age, PLoS ONE 13 (2018) e0201407. 10.1371/journal.pone.0201407.

[8] Z. Alam, N.K. Rendos, A.M. Vargas, J. Makanjuola, T.M. Kesar, Timing of propulsion-related biomechanical variables is impaired in individuals with post-stroke hemiparesis, Gait & Posture 96 (2022) 275–278. 10.1016/j.gaitpost.2022.05.022.

[9] A.D. Kuo, Energetics of actively powered locomotion using the simplest walking model, J Biomech Eng 124 (2002) 113–120. 10.1115/1.1427703.

[10] J.M. Donelan, R. Kram, A.D. Kuo, Simultaneous positive and negative external mechanical work in human walking, J Biomech 35 (2002) 117–124. 10.1016/s0021-9290(01)00169-5.

[11] O. Darici, A.D. Kuo, Humans optimally anticipate and compensate for an uneven step during walking, eLife 11 (2022) e65402. 10.7554/eLife.65402.

[12] S.S. Hosseini-Yazdi, Optimum Push-off During Uneven Walking for Just-in-Time Strategy; Delayed Push-off Exertion is Mechanically Costly, (2024). 10.1101/2024.07.14.603455.

[13] S.-S. Hosseini-Yazdi, J.E.A. Bertram, Center of mass work analysis predicts preferred walking speeds for varying walking conditions, Journal of Biomechanics 185 (2025) 112682. 10.1016/j.jbiomech.2025.112682.

[14] R.McN. Alexander, Simple Models of Human Movement, Applied Mechanics Reviews 48 (1995) 461–470. 10.1115/1.3005107.

[15] R.M. Alexander, Simple models of walking and jumping, Human Movement Science 11 (1992) 3–9. 10.1016/0167-9457(92)90045-D.

[16] A.D. Kuo, J.M. Donelan, A. Ruina, Energetic Consequences of Walking Like an Inverted Pendulum: Step-to-Step Transitions:, Exercise and Sport Sciences Reviews 33 (2005) 88– 97. 10.1097/00003677-200504000-00006.

[17] S.-S. Hosseini-Yazdi, J.E.A. Bertram, Optimum Push-off for Uneven Walking Based on the Just-in-Time Strategy: Walking with Interrupted Push-Off is Mechanically Costly, Journal of Biomechanical Engineering (2025) 1–20. 10.1115/1.4069666.

[18] S.-S. Hosseini-Yazdi, A.D. Kuo, The energetic cost of human walking as a function of uneven terrain amplitude, Journal of Experimental Biology (2025) jeb.249840. 10.1242/jeb.249840.

[19] M.H. Schwartz, A. Rozumalski, J.P. Trost, The effect of walking speed on the gait of typically developing children, J Biomech 41 (2008) 1639–1650. 10.1016/j.jbiomech.2008.03.015.

[20] Hosseini-Yazdi Seyed-Saleh, Energetics and Biomechanics of Uneven Walking for Young and Older Adults, University of Calgary, 2024. https://prism.ucalgary.ca/server/api/core/bitstreams/be2fa66b-26a2-4421-9d30-dfcc55dcfb35/content.

[21] O. Darici, H. Temeltas, A.D. Kuo, Optimal regulation of bipedal walking speed despite an unexpected bump in the road, PLoS ONE 13 (2018) e0204205. 10.1371/journal.pone.0204205.

[22] G.A. Cavagna, N.C. Heglund, C.R. Taylor, Mechanical work in terrestrial locomotion: two basic mechanisms for minimizing energy expenditure, Am J Physiol 233 (1977) R243–261. 10.1152/ajpregu.1977.233.5.R243.

[23] G.A. Cavagna, H. Thys, A. Zamboni, The sources of external work in level walking and running, J Physiol 262 (1976) 639–657. 10.1113/jphysiol.1976.sp011613.

[24] T.S. Keller, A.M. Weisberger, J.L. Ray, S.S. Hasan, R.G. Shiavi, D.M. Spengler, Relationship between vertical ground reaction force and speed during walking, slow jogging, and running, Clin Biomech (Bristol, Avon) 11 (1996) 253–259. 10.1016/0268-0033(95)00068-2.

[25] R.R. Neptune, S.A. Kautz, F.E. Zajac, Contributions of the individual ankle plantar flexors to support, forward progression and swing initiation during walking, Journal of Biomechanics 34 (2001) 1387–1398. 10.1016/s0021-9290(01)00105-1.

[26] A.H. Dewolf, Y.P. Ivanenko, F. Lacquaniti, P.A. Willems, Pendular energy transduction within the step during human walking on slopes at different speeds, PLOS ONE 12 (2017) e0186963. 10.1371/journal.pone.0186963.

[27] S.-S. Hosseini-Yazdi, J.E. Bertram, The consequence of uneven walking transitory modulation strategies: A simulation-based approach, J Theor Biol 614 (2025) 112234. 10.1016/j.jtbi.2025.112234.

[28] S.-S. Hosseini-Yazdi, A.D. Kuo, A split-belt instrumented treadmill with uneven terrain, Journal of Biomechanics 176 (2024) 112376. 10.1016/j.jbiomech.2024.112376.

[29] L.N. Awad, M.D. Lewek, T.M. Kesar, J.R. Franz, M.G. Bowden, These legs were made for propulsion: advancing the diagnosis and treatment of post-stroke propulsion deficits, J NeuroEngineering Rehabil 17 (2020) 139. 10.1186/s12984-020-00747-6.

[30] D.J. Farris, A. Hampton, M.D. Lewek, G.S. Sawicki, Revisiting the mechanics and energetics of walking in individuals with chronic hemiparesis following stroke: from individual limbs to lower limb joints, J NeuroEngineering Rehabil 12 (2015) 24. 10.1186/s12984-015-0012-x.

[31] T.J. Van Der Zee, A.D. Kuo, Soft tissue deformations explain most of the mechanical work variations of human walking, Journal of Experimental Biology 224 (2021) jeb239889. 10.1242/jeb.239889.

[32] K.E. Zelik, A.D. Kuo, Human walking isn’t all hard work: evidence of soft tissue contributions to energy dissipation and return, Journal of Experimental Biology 213 (2010) 4257–4264. 10.1242/jeb.044297.

[33] D.S. Rooks, D.P. Kiel, C. Parsons, W.C. Hayes, Self-Paced Resistance Training and Walking Exercise in Community-Dwelling Older Adults: Effects on Neuromotor Performance, J Gerontol A Biol Sci Med Sci 52A (1997) M161–M168. 10.1093/gerona/52A.3.M161.

